# Diversity of hydrodynamic radii of intrinsically disordered proteins

**DOI:** 10.1101/2023.06.12.543592

**Authors:** Michał K. Białobrzewski, Barbara P. Klepka, Agnieszka Michaś, Maja K. Cieplak-Rotowska, Zuzanna Staszałek, Anna Niedźwiecka

## Abstract

Intrinsically disordered proteins (IDPs) form an important class of biomolecules regulating biological processes in higher organisms. The lack of a fixed spatial structure facilitates them to perform their regulatory functions. Due to the possibility of large conformational changes of IDPs, the cellular milieu can also control productivity of biochemical reactions. From the biophysical point of view, IDPs are biopolymers with a broad configuration state space. The conformation of such a biopolymer depends on non-covalent interactions of its amino acid side chain groups at given temperature and chemical conditions. Thus, the hydrodynamic radius (R_h_) of an IDP of a given polymer length (N) is a sequence- and environment-dependent variable. We have reviewed the literature values of hydrodynamic radii of IDPs determined experimentally by SEC, AUC, PFG NMR, DLS, and FCS, and complement them with our FCS results obtained for a series of protein fragments involved in regulation of human gene expression. The data collected herein show that the values of hydrodynamic radii of intrinsically disordered proteins can span the full space between the folded globular and denatured proteins in the R_h_(N) diagram.

## Introduction

Intrinsically disordered proteins (IDPs) are nowadays recognized as an important class of biomolecular players (Dyson and Wright 2005). In eukaryotes, bioinformatics predictions suggest that *ca.* one third of the proteome could contain intrinsically disordered regions (IDRs) (Ward et al. 2004). The IDPs have been even called “the dark matter of molecular biology” (Darling and Uversky 2018) due to the significant roles they play in a living cell and the challenges associated with limited possibilities of their detection and systematic study. The main drawback in the traditional way of thinking about the structure-function relationship in the context of IDPs is the lack of a constant, stable, well-defined three-dimensional structure that could be resolved by X-ray crystallography, multidimensional nuclear magnetic resonance or cryo-electron microscopy and shed light on the structural requirements for their activity. In contrast, it is the lack of a fixed spatial structure that is crucial to the functions that IDPs perform, either by providing conformational flexibility to multidomain proteins or by direct interactions of their IDRs encompassing short linear motifs (SLiMs) (Bhandari et al. 2014). IDPs are usually involved in superior cellular regulatory processes, such as *e.g.* transcription regulation (Borgia et al. 2018), mRNA maturation (Loughlin and Wilce 2019), post-transcriptional gene silencing (Sheu-Gruttadauria and MacRae 2018), translation initiation regulation (Fletcher et al. 1998; Gingras et al. 2001), transmembrane signalling (Seiffert et al. 2020) or biomineralization (Evans 2019). In particular, many proteins implicated in mRNA turnover that constitute the cytoplasmic membrane-less organelles (MLOs) called RNA processing bodies (P-bodies) are IDPs (Banani et al. 2016; Nosella and Forman-Kay 2021; Currie et al. 2023). Cellular MLOs are biomolecular condensates that are thought to emerge as a result of spontaneous liquid-liquid phase separation (LLPS) and further maturation of the IDP-rich, denser phase, based on the polymeric character of the IDPs and their multivalency (Brangwynne et al. 2015; Uversky 2017; Alberti et al. 2019; Abyzov et al. 2022). The IDPs thus contribute to membrane-less compartmentalization of the cellular processes that need to be separated in space and time (O’Flynn and Mittag 2021; Musacchio 2022). Consequently, IDPs are often associated with cancer related processes and other diseases (Babu et al. 2011; Krois et al. 2018).

Among MLOs, one can distinguish GW-bodies as a subgroup of P-bodies (Eystathioy et al. 2002; Jakymiw et al. 2007). They are abundant in GW182, a glycine- and tryptophan-rich protein of 182 kDa mass (Eystathioy et al. 2002), involved in microRNA-dependent gene silencing. GW182 is responsible for linking the microRNA-targeted mRNA with the CCR4-NOT deadenylase complex by SLIMs interactions (Fabian et al. 2011; Braun et al. 2013; Mathys et al. 2014). The C-terminal silencing domain (SD) of GW182 was shown experimentally by hydrogen-deuterium exchange mass spectrometry to be intrinsically disordered (Cieplak-Rotowska et al. 2018). The GW182 SD interacts *i.a.* with a middle domain (M) of a huge scaffolding CNOT1 subunit (2 300 amino acid residues) of CCR4-NOT (Fabian et al. 2011). While CNOT1 is essentially α-helical, it contains also long loops linking the helices; therefore, their different fragments can be either folded into a stable 3D structure or intrinsically disordered. Besides CCR4-NOT that serves for regular mRNA surveillance, there is another 3’ poly(A)-specific nuclease (PARN) that plays a crucial role *i.a.* in maternal mRNA removal during early development and telomerase RNA component deadenylation (Tummala et al. 2015; reviewed in: Virtanen et al. 2013). PARN forms a covalently bound homodimer, where each monomer encompasses three folded domains (nuclease, R3H and RRM) (Wu et al. 2005, 2009) and an over 100 amino acid residues long C-terminal intrinsically disordered tail (PARN C) (Niedzwiecka et al. 2011) that contains a nuclear localization signal. Here, we analyse hydrodynamic properties of various protein fragments of human GW182 SD, CNOT1 M and PARN C (Fig. 1, Supplementary Information).

**Fig. 1.**
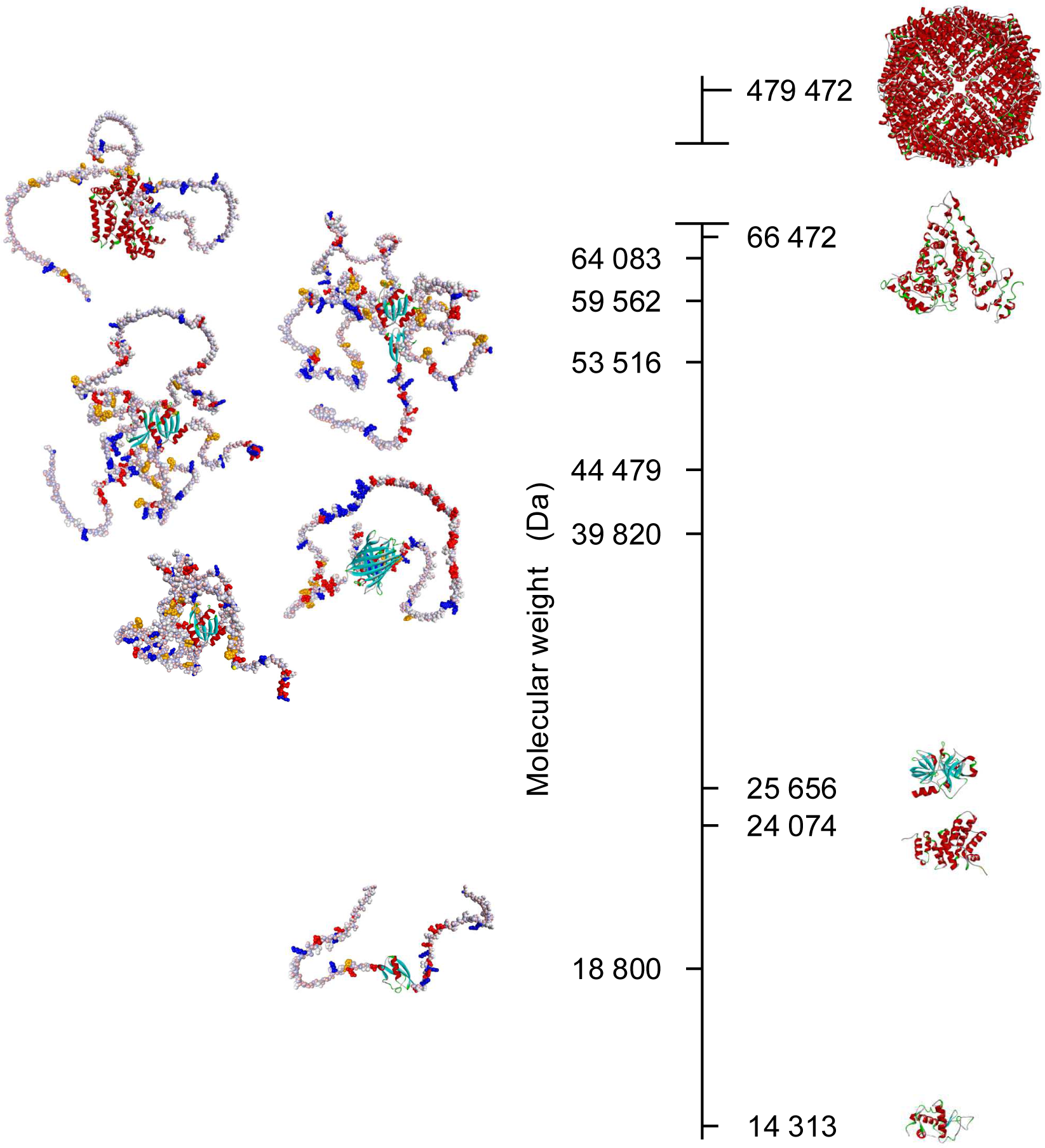
Example putative conformations of the intrinsically disordered proteins studied in this work (left panel) ordered top-down according to their molecular weights, CNOT1 M long, SUMO-GW182 SD, SUMO-GW182 SD10, PARN C-mCherry, GW182 SD10, SUMO-GW182 SD peptide, predicted by AlphaFold 2 (Jumper et al. 2021; Mirdita et al. 2022) in comparison with the globular proteins (right panel), apoferritin (PDB id. code: 7vd8) (Fan et al. 2021), human serum albumin (PDB id. code: 1uor) (He and Carter 1992), α-chymotrypsinogen A (PDB id. code: 2cga) (Wang et al. 1985), CNOT1 M short (CNOT1(800-999), PDB id. code: 4j8s) (Fabian et al. 2013), and lysozyme (PDB id. code: 1e8l) (Schwalbe et al. 2001). The N-terminal fragment lacking in the crystal structure of CNOT1 M short was generated by AlphaFold 2. Intrinsically disordered protein fragments are shown in the CPK sphere atom representation coloured according to partial charges, except for positively charged amino acid residues (Arg, Lys) marked in blue, negatively charged residues (Asp, Glu) in red, and hydrophobic aromatic residues (Trp, Phe) in dark yellow. Folded fragments are shown as ribbon coloured according to secondary structural elements, α-helices in red, β-sheets in cyan, and β-turns in green

IDPs are biopolymers with a broad and almost continuous configuration state space. They can change their conformations to a great extent under the influence of physicochemical conditions (Uversky 2009; Moses et al. 2020). This means that due to the possibility of significant conformational changes, environmental conditions may affect the accessibility of the IDP binding sites. A spectacular example is related to the eukaryotic translation initiation inhibitors, eIF4E-binding proteins (4E-BPs) (Pause et al. 1994). In the pioneering works to the field of IDPs, the NMR studies of 4E-BP1 revealed that the active protein is unstructured (Fletcher et al. 1998; Fletcher and Wagner 1998). It was also shown that the multistage, hierarchical phosphorylation of 4E-BP1 is necessary to trigger its dissociation from eIF4E (Gingras et al. 2001; Niedzwiecka et al. 2002). The initial biophysical explanation of the affinity loss of the hyperphosphorylated 4E-BP was based on electrostatic repulsion, since the 4E-BPs-binding site of eIF4E is negatively charged (Marcotrigiano et al. 1999). However, further studies of a homologous 4E-BP2 proved that the phosphorylation changed the hydrogen bonding possibilities, leading to formation of a 3D folded structure that sequestered a tyrosine residue that was crucial for the eIF4E binding (Bah et al. 2015).

Because of the thermodynamic driving forces of conformational changes of IDPs, their hydrodynamic radius (*R_h_*) is also temperature dependent in a non-trivial way, *e.g.* the *R_h_* of a p53 protein fragment (residues 1-93) was shown to even decrease with increasing temperature due to the heat-induced compaction of the IDP structure (Langridge et al. 2014). Thus, the *R_h_* of an IDP is a variable and not a constant value describing the hydrodynamic properties of a biomolecule, contrary to *e.g.* chemical dyes with a fixed spatial structure. Consequently, the diffusion coefficient (*D*) of an IDP depends on all the chemical factors that can influence the molecule dimensions (Moses et al. 2020) and on temperature (*T*) in a more intricate way through *R_h_*:

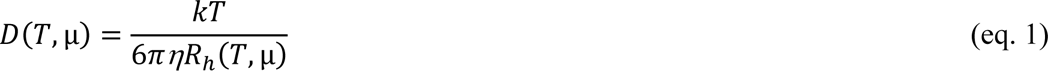

where *µ* is the chemical potential reflecting *i.a.* the pH value, ionic strength, osmotic stress and the presence of all putatively interacting small molecules or ions. The conformational heterogeneity of an IDP can thus lead to a range of values of its hydrodynamic parameters at given temperature, instead of a single value. The cellular and extracellular milieux may determine this way the kinetics of diffusion-controlled intermolecular reactions involving IDPs *via* their conformational changes.

In general, a wide range of IDP conformational variability determined by different environmental factors can be a kind of switch regulating the effectiveness of the processes occurring at the molecular level, *e.g.* formation of protein-protein or protein-nucleic acid complexes or emergence of larger aggregates and microcrystals.

According to the well-established theory of diffusion of polymers (Flory 1949; Le Guillou and Zinn-Justin 1977), *R_h_* can be approximated by a power function of the number of the polymer units (*N*):

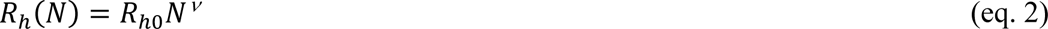

where *ν* is the critical exponent equal to 1/3 for polymers perfectly folded into spheres and *ν* = 3/5 for unfolded linear polymers. Within this formalism, *N* is ascribed to the number of amino acid residues in a protein chain on the assumption that they are indistinguishable. The hydrodynamic radii measured for both folded globular proteins and denatured proteins seem to follow the polymer theory with the above critical exponent values, respectively (Marsh and Forman-Kay 2010). However, it was a matter of debate whether it would be possible to determine a specific value of the critical exponent for the whole IDP class (Le Guillou and Zinn-Justin 1977) or predict the *R_h_* value for a known IDP sequence without employing molecular dynamics simulations or recalculation from the radius of gyration (*R_g_*) measured by other experimental approaches (Marsh and Forman-Kay 2010).

In order to gain a better understanding of the complex nature of hydrodynamic properties of IDPs, we have gathered an exhaustive set of available experimental literature results for IDPs’ hydrodynamic radii determined by size excluded chromatography (SEC), analytical ultracentrifugation (AUC), pulsed field gradient NMR (PFG NMR), dynamic light scattering (DLS), and fluorescence correlation spectroscopy (FCS). Against the background of the literature data, we present our FCS results obtained for a series of protein fragments involved in regulation of human gene expression. The experimental data collected herein show unambiguously that the values of hydrodynamic radii of intrinsically disordered proteins span the full space between the folded globular and denatured proteins in the *R_h_*(*N*) diagram.

## Materials and Methods

### Review of literature data

The experimental data were selected from literature based on the following criteria:

1. the *R_h_* values have been determined directly from appropriate experiments, without conversions from other experimental quantities such as *R_g_* that would require some assumptions;

2. the *R_h_* values could be ascribed to the protein sequences found in the literature or in the UniProtKB database.

The data of the selected proteins are gathered in Supplementary information (Hemmings et al. 1984; McCubbin et al. 1985; Lynch et al. 1987; Donaldson and Capone 1992; Uversky et al. 1999, 2002b, a; Wilkins et al. 1999; Guez et al. 2000; Campbell et al. 2000; Bouvier and Stafford 2000; Adkins and Lumb 2002; Karlin et al. 2002; Denning et al. 2002, 2003; Permyakov et al. 2003; Zeev-Ben-Mordehai et al. 2003; Tcherkasskaya et al. 2003; Longhi et al. 2003; Mayor et al. 2003; Chong et al. 2004; Sánchez-Puig et al. 2005a, b; Yiu et al. 2006; Goldgur et al. 2007; Geething and Spudich 2007; Magidovich et al. 2007; Khaymina et al. 2007; Gall et al. 2007; Yi et al. 2007; Lowry et al. 2008; Zhang et al. 2008; Paleologou et al. 2008; Kapłon et al. 2008; Krishnan et al. 2008; Paz et al. 2008; Soragni et al. 2008; Haaning et al. 2008; Sivakolundu et al. 2008; Danielsson et al. 2008; Baker 2009; Neira et al. 2009; Habchi et al. 2010; Choi et al. 2011; Nag et al. 2011; Perez et al. 2014; Langridge et al. 2014; Poznar et al. 2017; Wollenhaupt et al. 2018; Więch et al. 2019; Chatterjee and Pollard 2019).

### Chemicals

The chemicals for protein expression and purification were purchased from Merck-Sigma-Aldrich and were analytically pure (grade A). The AF488 NHS ester was purchased from Lumiprobe GmbH. Alexa Fluor 546 NHS ester was purchased from Invitrogen.

### Protein expression, purification and labelling

The protein sequences are given in the Supplementary information. CNOT1 M long, CNOT1 M short, SUMO-GW182 SD, SUMO-GW182 SD10, GW182 SD10 were expressed and purified as described previously (Fabian et al. 2011, 2013; Cieplak-Rotowska et al. 2018). The genes for SUMO-GW182 SD peptide and PARN C-mCherry protein constructs were ordered from BioCat GmbH (Heidelberg, Germany) in the form of His_6_-SUMO fusions. The proteins were overexpressed in *Escherichia coli* Rosetta 2(DE3)pLysS, purified in the form of fusion proteins by Ni-NTA affinity purification and further digested by using the SUMO Protease (Sigma-Aldrich) according to the protocols provided by the manufacturer. The proteins were further purified by anion exchange (HiTrap Q, Cytiva) or SEC (Superdex 200 Increase 10/300 GL, Cytiva) chromatography at ÄKTA FPLC (system ÄKTA pure; GE Healthcare). The purity of the protein was checked by the SDS PAGE.

Proteins were labelled by using the AF488-NHS ester according to the manufacturer’s protocol and purified from the excess of the dye by Zeba spin columns (Thermo Scientific), dialysis (Pur-A-Lyzer, Sigma-Aldrich) or SEC (Superdex 200 Increase 10/300 GL, Cytiva), depending on the protein properties.

### Fluorescence correlation spectroscopy measurements

The FCS experiments were done at Zeiss LSM 780 with ConfoCor 3, in 50 mM Tris/HCl buffer pH 8.0, 150 mM NaCl, 0.5 mM EDTA, and 1 mM TCEP, in droplets of 25 µl, at 25 ± 0.5 °C. The temperature was measured inside the droplet after the FCS measurements by means of a micro-thermocouple. The buffer and samples were filtered through 0.22 μm before the experiment. The solution viscosity was determined by comparison of the AF488 diffusion time in pure water (*D_AF488_* = 435 μm^2^ s^−1^) (Petrásek and Schwille 2008) and in the buffer at the same equipment calibration. The structural parameter (*s*) was determined every time with use of AF488 or AF546 (*D* = 341 μm^2^ s^−1^) (Petrásek and Schwille 2008), individually for each microscopic slide previously passivated with BSA in the buffer.

The experiments for proteins labelled by AF488 were performed at the 488 nm excitation wavelength with a relative Argon multiline laser power of 3 %, MBS 488 nm, BP 495-555 nm. A single measurement time was 3 to 6 s, repeated 10 to 50 times in a set. The set of measurements was repeated 3 times in 5 independent droplets.

For AF546 and the mCherry-fused protein, the excitation wavelength was 561 nm at 2 % relative DPSS laser power, MBS 488/561 nm, LP 580 nm. A single measurement time was 3 s, repeated 50 times in a set. The set was repeated 3 times in 5 independent droplets. A dampening factor of 10 % and a dust filter of 10 % were applied.

Photophysical processes of AF488 and mCherry were investigated in independent sets of experiments. A relative laser power ranging from 3 to 20 % at 488 nm was used for AF488. The average triplet state lifetime of AF488 was about 4 µs. In the case of mCherry, the measurements were performed in 30 % glycerol to slow down the protein diffusion and extract the blinking (Hendrix et al. 2008), at a 7 % relative laser power. The fraction of mCherry population that undergoes blinking was found to be about 24 % both for mCherry alone and in the fusion constructs with the GW182 SD sequences.

### Data analysis

The FCS raw data were analysed by Zen2010 (Zeiss) by global fitting of the autocorrelation curve to a set of 10 to 50 single measurements, after detailed inspection of each measurements in order to exclude possible oligomerization or aggregation of the sample in the confocal volume during the experiment. The autocorrelation function providing for 3D diffusion and photophysical processes was fitted according to the equations (Sahoo and Schwille 2011):

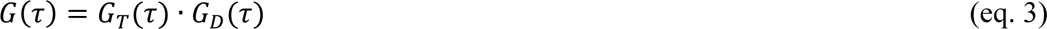

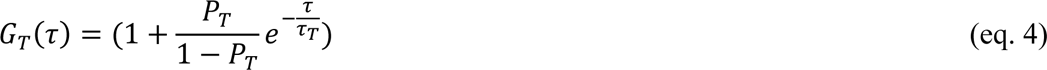

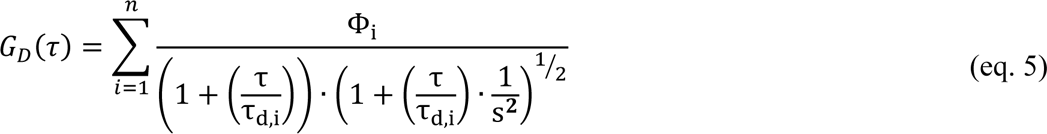

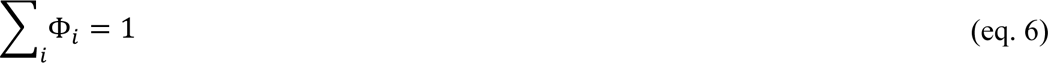

where: *G*(τ) is the fitted autocorrelation function; *G_T_*(τ), normalized autocorrelation function for photophysical processes; *G_D_*(τ), autocorrelation function for the diffusion of *n* components; *P_T_*, triplet state or blinking fraction; τ_T_, lifetime of the photophysical process; τ_*d,i*_, diffusion time for the *i*-th component; *s*, structural parameter of the confocal volume.

A one-component model (*n* = 1) providing for mCherry blinking was fitted for the fusion fluorescent proteins, and a two-component model (*n* = 2) taking into account the AF488 triplet state and the presence of a residual freely diffusing dye was used for the chemically labelled proteins. The fraction of free AF488 in the experiment with HSA was ∼14 %

The mCherry blinking fraction and the AF488 triplet state lifetime were determined from the independent experiments and fixed during the global analysis.

The protein *R_h_*was determined from the diffusion time, τ_*d*_, providing for the actual buffer viscosity

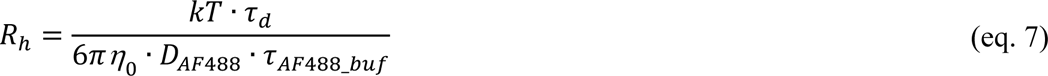

where *η_0_* is the viscosity of pure water (IAPWS 2008) at the temperature *T* and τ_*AF*488_*buf*_ is the AF488 diffusion time in the buffer at the same calibration.

Other non-linear or linear numerical regressions were performed by Prism 6 (GraphPad Software, Inc.).

The total experimental uncertainty was determined according to the rules for propagation of small errors (Taylor 1997), taking into account both numerical uncertainty of the fitting, statistical dispersion of the results and uncertainties of other experimental values used for calculation of the results.

### Bioinformatics

Example conformations of the IDPs were generated by AlphaFold 2.0 notebook (Jumper et al. 2021; Mirdita et al. 2022). Protein structures were drawn by using Discovery Studio v3.5 (Accelrys Software, Inc.).

## Results and Discussion

Among many experimental techniques that can yield the *in vitro* diffusion coefficients, the FCS method is the only one that measures the true self-diffusion coefficient of a molecule due to the possibility of the free 3D diffusion in a droplet and negligible protein-protein interactions at nanomolar concentrations. DLS and PFG NMR need higher protein concentrations, SEC is based on a spatially limited diffusion through pores in a chromatographic bed in the flow under pressure, and AUC measures diffusion in experiments with an applied high centrifugal force. Moreover, even the choice of a data analysis formalism can change the final outcome (Petsev et al. 2000). For these reasons, the results obtained for the same protein at similar temperature and solution conditions by different hydrodynamic methods do not necessarily match exactly (Karlin et al. 2002; Longhi et al. 2003; Geething and Spudich 2007; Paz et al. 2008; Habchi et al. 2010; Perez et al. 2014; Poznar et al. 2017). Nevertheless, these values converge usually within the limit of 3 σ and therefore it seems interesting to collect here all available *bona fide* experimental results for hydrodynamic radii of IDPs from the scientific literature, together with the FCS results measured for some of our protein constructs and compare them with the extremes for folded globular and denatured proteins.

We have determined *R_h_*for several protein constructs related to regulation of human gene expression. Most of these proteins are experimentally proven (Cieplak-Rotowska et al. 2018) or bioinformatically predicted (Niedzwiecka et al. 2011) to be IDPs. Their times of diffusion through the confocal volume of the microscope are much larger than for a hypothetical protein of the same mass or the polypeptide chain length but packed into a spherical shape. An example FCS autocorrelation curve for an intrinsically disordered C-terminal fragment of the poly(A)-specific ribonuclease (PARN C) fused by a flexible linker with a fluorescent mCherry protein is presented in Fig. 2 together with the curve for a folded protein, human serum albumin (HSA), being a reference standard used commonly for measurements of protein molecular masses and radii. It is clear that the IDP having a shorter polypeptide chain and a lower molecular weight (395 residues, 44 479 Da) is characterised by a longer diffusion time compared to the folded globular protein of a longer polypeptide chain and of a larger molecular weight (585 residues, 66 472 Da). Consequently, the *R_h_* value of the intrinsically disordered PARN C significantly exceeds the value that could be ascribed to the corresponding folded globular protein.

**Fig. 2.**
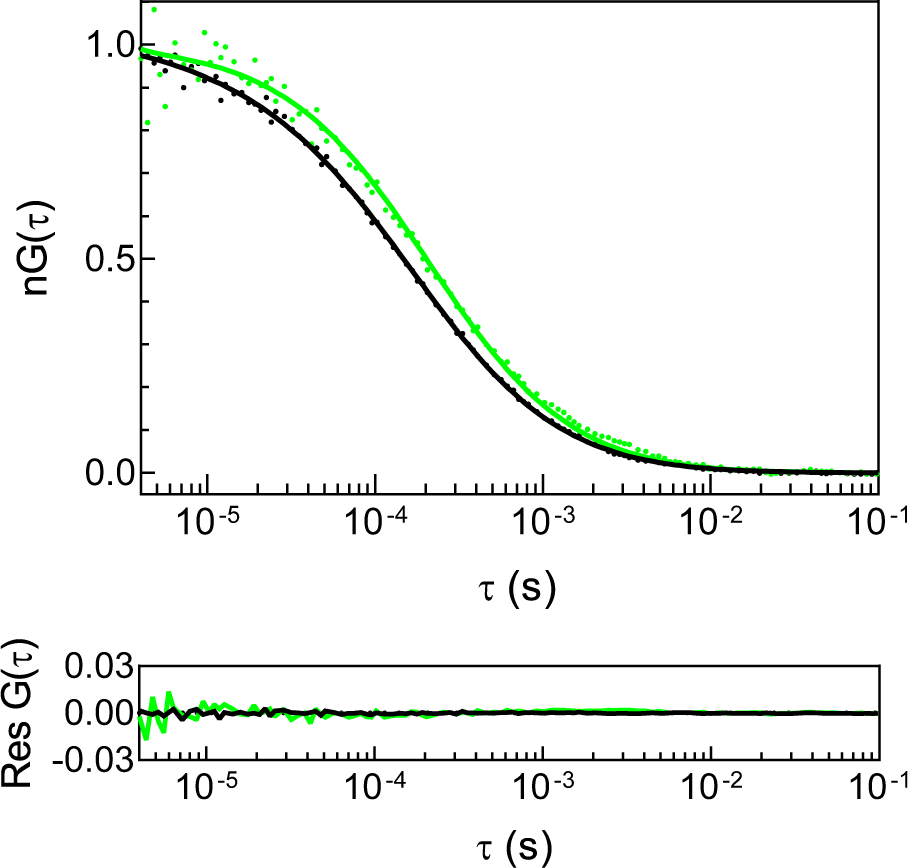
Example normalized FCS autocorrelation curves with non-normalized fitting residuals for a folded protein, HSA (black), and an intrinsically disordered protein, PARN C-mCherry (green), in 50 mM Tris/HCl buffer pH 8.0, 150 mM NaCl, 0.5 mM EDTA, 1 mM TCEP, at 298 K. Dots, experimental points; lines, curves fitted according to eq. 3

The FCS results obtained for the IDPs and globular proteins in this study are gathered in Table 1 and shown in Fig. 3. Contrary to the globular CNOT1 M short fragment known from the crystal structure (Fabian et al. 2013), the intrinsically disordered CNOT1 M long, GW182 SD and PARN C protein fragments, encompassing short or long disordered regions, outlie the 95% confidence interval of the straight line in Fig. 3a and the curve in Fig 3b for the standard folded globular proteins. Interestingly, the *R_h_* values for the GW182 SD constructs may suggest that even if the protein sequence is dominated by IDRs, the *R_h_* value of the IDP can be closer to the value expected for a folded globular protein of the same molecular weight or length than to the denatured protein, which is in contrast to PARN C. This diverse behaviour can be attributed to different sequence biases in the case of GW182 SD and PARN C. While the PARN C sequence represents a block-like structure comprising a bipartite positively charged nuclear localization signal separated by a negatively charged region, GW182 SD resembles a polyampholyte polymer with an alternating charge pattern and is especially rich in water-exposed tryptophan side chains next to glycine residues. Consequently, GW182 SD could form a kind of entropy-driven, loose and dynamic hydrophobic quasi-core due to the presence of the tryptophan residues, instead of a random coil. This possibility is also suggested by the AlphaFold 2.0 (Jumper et al. 2021; Mirdita et al. 2022) prediction of a putative GW182 SD conformation (Fig. 1), although it should be treated with caution and need future molecular dynamics simulations within a proper model for such a long IDP chain.

**Fig. 3.**
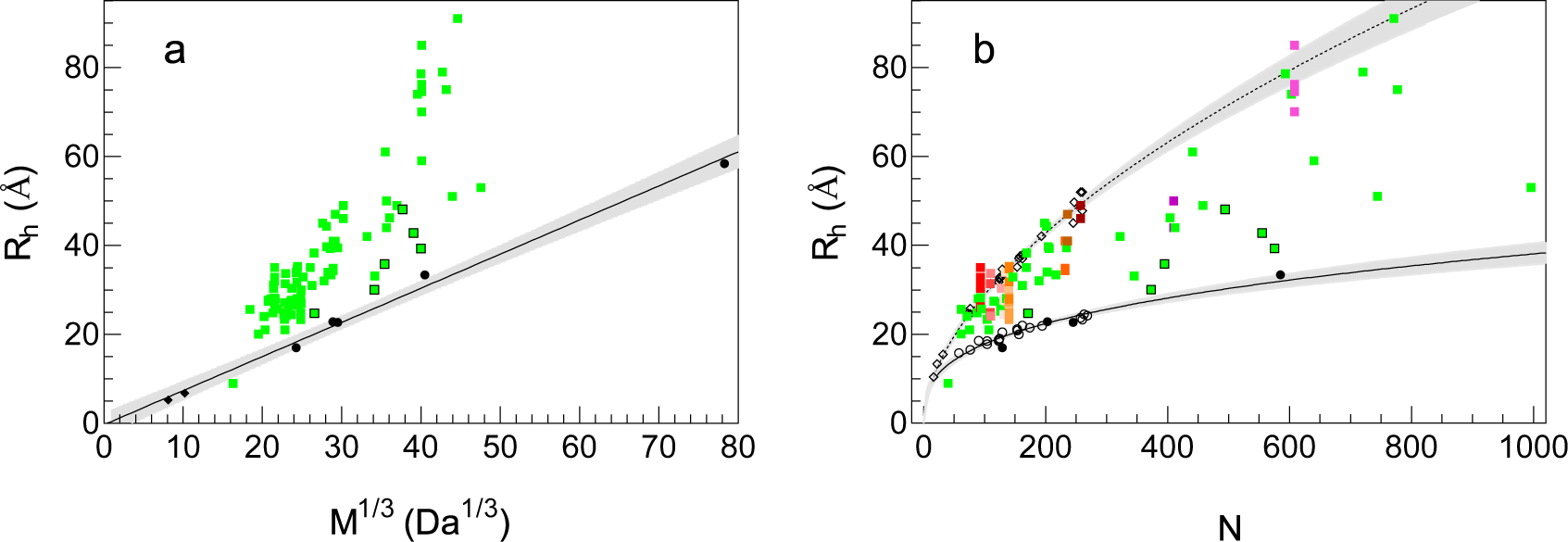
Hydrodynamic radii, *R_h_*, of intrinsically disordered proteins (black bordered green squares), globular proteins (black circles) and chemical fluorescent probes (AF488 and Alexa Fluor 546, black diamonds) determined by FCS in this study (Table 1) (a) as a function of the molecular mass, *M*, compared to literature *R_h_* values for IDPs (plain green squares, Table S1); and (b) as a function of the polymer residue number, *N*, compared to literature data (plain squares, Table S1), where the results for the same protein at different conditions are marked in shades of red-orange-magenta; open circles and diamonds denote folded globular and chemically denatured proteins from literature, respectively. Chemical fluorescent probes and apoferritin are not taken into account in (b). The analytical curve according to eq. 2 was fitted for folded proteins to literature data merged with the results of this study, and for denatured proteins to literature data. The grey shaded regions represent the 95% confidence interval bands

**Table 1.**
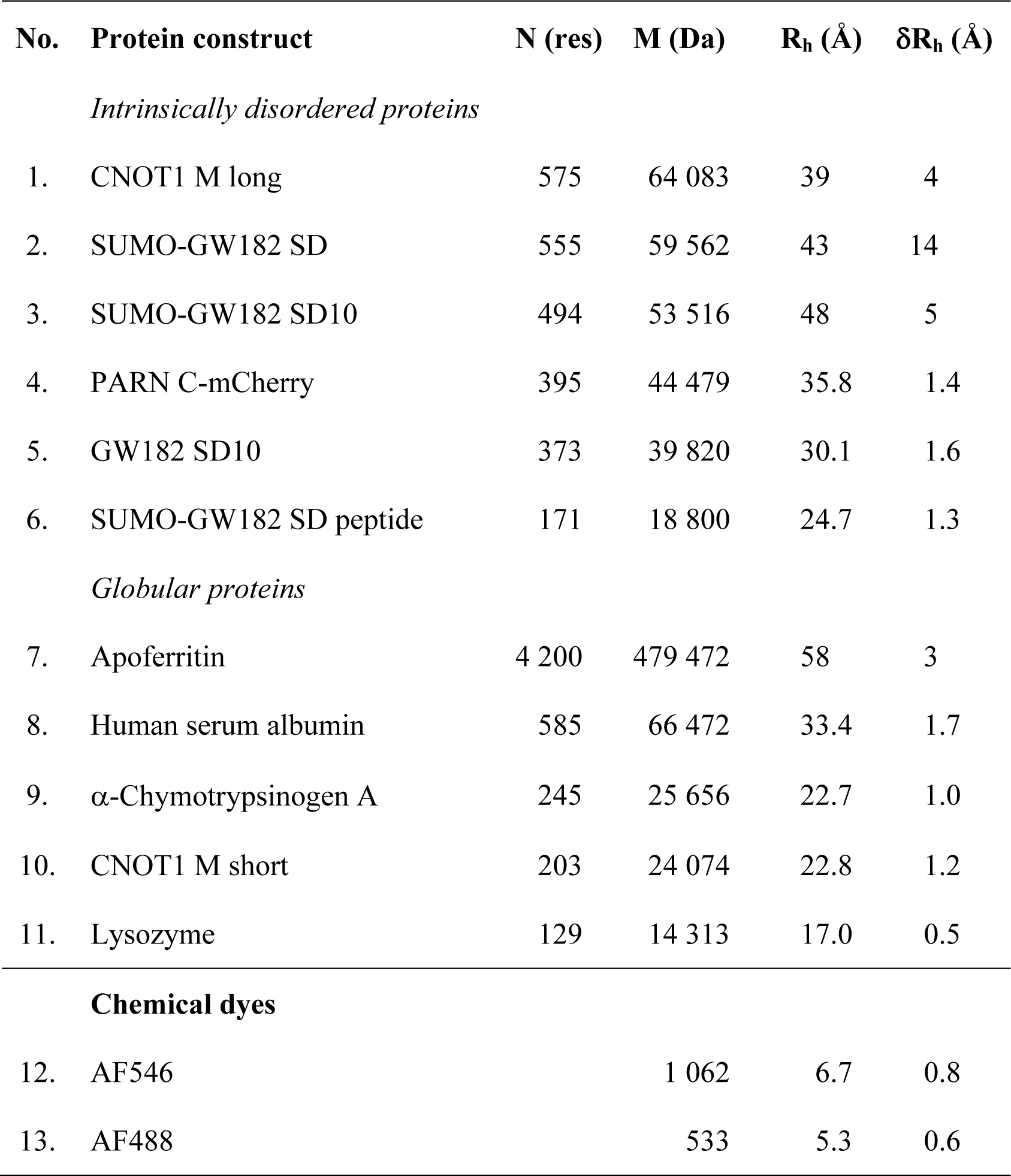
Hydrodynamic radii, *R_h_*, of protein constructs and chemical dyes determined in this work by fluorescence correlation spectroscopy, in 50 mM Tris/HCl buffer pH 8.0, 150 mM NaCl, 0.5 mM EDTA, and 1 mM TCEP, at 25 °C. *N*, number of amino acid residues; *M,* molecular weight of the protein construct, *δR_h_*, experimental uncertainty of *R_h_*

A collection of a broad range of experimentally determined hydrodynamic radii of IDPs from literature (Supplementary Table S1) confirms that the shape of an individual protein can differ from the ideal sphere to a various extent (Fig. 3a). This is visible as a larger or smaller deviation of the *R_h_* value of the IDPs from the straight line describing *R_h_* as a linear function of the cube root of the molecular weight, which is quite well applicable to the folded globular proteins and small molecules (slope 0.77 ± 0.03 Å·mol/g, intercept -0.32 ± 1.05 Å).

A more comprehensive analysis can be made, however, based on the polymer model referring to the number of the chain units (*N*), *i.e.* the polypeptide amino acid residues. This approach enables us to analyse the hydrodynamic behaviour of protein conformations influenced not only by pH, electrostatic screening by an ionic strength or the presence of an unspecific chemical denaturant but also by the changes that modify the effective molecular mass of the protein and keep the *N* constant. Those are point mutations, posttranslational modifications and stoichiometric binding of ions or small molecules at different environmental conditions. In the set of the *R_h_* values of IDPs shown in Fig. 3b, the variants of the same proteins are marked in the shades of red-orange-magenta. The Fig. 3b makes it evident that the changes in the *R_h_* values can be very large even for the same IDP.

The issue of the diverse dimensions of the same IDP chain at different solution conditions has been elaborated in much detail based on radii of gyration (*R_g_*) of IDPs measured by single molecule Förster resonance energy transfer (smFRET) (Hofmann et al. 2012). Highly charged proteins can first undergo a tightening (an *R_g_* decrease) and only then an expansion (an *R_g_* increase) of their chain with the increasing concentration of an ionic denaturant due to the interplay of the electrostatic screening and the denaturant binding (Müller-Späth et al. 2010). A similar effect was also observed in bulk SEC measurements of the *R_h_* values of a highly acidic protein involved in biomineralization (Poznar et al. 2017). This is hence not surprising that IDPs do not form a uniform class of proteins that could be assigned a single critical exponent value in the polymer model. To sum up, due to the diversity of the conformations acquired by different IDPs dependent on their amino acid side chain properties and environmental conditions, the *R_h_*values of IDPs can adopt any value between the limiting curves for *ν* = 1/3 and 3/5.

The critical exponent determined here for the *R_h_* values of the folded globular proteins gathered from literature (Wilkins et al. 1999; Tcherkasskaya et al. 2003) merged together with our data (Table 1) is *ν_f_* = 0.33 ± 0.03 (95 % CI), which is in agreement with the value of 1/3 for a polymer folded tightly into a spherical shape (Flory 1949). The value for the denatured proteins (Wilkins et al. 1999) is *ν_d_* = 0.56 ± 0.03 (95 % CI).

While the simple polymer model works well for estimation of hydrodynamic properties of both folded globular and completely unfolded proteins, the IDP properties are beyond the predicting capability of this model. Many approaches have been recently developed to address this issue *e.g.* (Rodríguez Schmidt et al. 2012; Mittal et al. 2013, 2018; Nygaard et al. 2017; Peti et al. 2018; Das et al. 2018; Estaña et al. 2019; Baul et al. 2019; Gomes et al. 2020; Pesce et al. 2023). However, their application needs either molecular dynamics simulations with a risk of a strong bias coming from the force field choice (Rauscher et al. 2015), an experimental input in the form of SAXS, NMR, FRET measurements, some assumptions enabling recalculation of *R_h_* from *R_g_*or is limited by the IDP lengths. An effective numerical method for fast prediction of the *R_h_*values for any large IDP at given conditions is still lacking, hence elaboration of new theoretical models such as *e.g.* (Cichocki et al. 2019) would be useful.

## Conclusions

The results obtained for the protein fragments related to the proteins involved in human gene silencing (GW182, CNOT1, PARN) determined by FCS, together with the collection of the literature experimental data show that hydrodynamic radii of IDPs of given lengths can acquire any value in between the power function curves describing folded globular and denatured proteins. New theoretical and semi-empirical models are necessary to enable fast estimation of hydrodynamic properties of large IDPs taking into account their various conformational states that depend on the sequence, post-translational modifications and the chemical milieu.

## Statements and Declarations

The authors have no relevant financial or non-financial interests to disclose

## Acknowledgements

The work was supported by the National Science Centre grant no. UMO-2016/22/E/NZ1/00656. The studies were performed in the NanoFun laboratories co-financed by the European Regional Development Fund within the POIG.02.02.00-00-025/09 Program. We thank Prof. Nahum Sonenberg and Dr. Marc Fabian for sharing plasmids for some protein constructs, Dr. Remigiusz Worch for helpful discussion, and Ms. Magdalena Duszka for excellent technical assistance.

