## Supplemental Information for "Diversity of hydrodynamic radii of intrinsically disordered proteins"

European Biophysics Journal

Michał K. Białobrzewski[1], Barbara P. Klepka[1], Agnieszka Michaś[1], Maja K. Cieplak-Rotowska[1,2,3], Zuzanna Staszałek[1], Anna Niedźwiecka[1,*]

[1] *Laboratory of Biological Physics, Institute of Physics, Polish Academy of Sciences, Aleja Lotnikow 32/46, PL-02668, Warsaw, Poland*

[2] *Division of Biophysics, Institute of Experimental Physics, Faculty of Physics, University of Warsaw, Pasteura 5, PL-02093, Warsaw, Poland*

[3] *present address: The International Institute of Molecular Mechanisms and Machines, Polish Academy of Sciences, Flisa 6, PL-02247 Warsaw, Poland*

**Data of protein constructs studied in this work**

Protein name, schematic sequence composition and full sequence

**CNOT1 M long**

linker-CNOT1(728-1267)-$His_6$

```
GPLGSPEFPG RPMASMTGGQ QMGGSGSGSP HTQSMQGFPP NLGSAFSTPQ SPAKAFPPLS
TPNQTTAFSG IGGLSSQLPV XGLGTGSLTG IGTGALGLPA VNNDPFVQRK LGTSGLNQPT
FQQTDLSQVW PEANQHFSKE IDDEANSYFQ RIYNHPPHPT MSVDEVLEML XRFKDSTIKR
EREVFNCMLR NLFEEYRFFP QYPDKELHIT ACLFGGIIEK GLVTYMALGL ALRYVLEALR
KPFGSKMYYF GIAALDRFKN RLKDYPQYCQ HLASISHFMQ FPHHLQEYIE YGQQSRDPPV
KMQGSITTPG SIALAQAQAQ AQVPAKAPLA GQVSTMVTTS TTTTVAKTVT VTRPTGVSFK
KDVPPSINTT NIDTLLVATD QTERIVEPPE NIQEKIAFIF NNLSQSNMTQ KVEELKETVK
EEFMPWVSQY LVMKRVSIEP NFHSLYSNFL DTLKNPEFNK MVXNETYRNI KVLLTXDKAA
ANFSDRSLLK NLGHWLGMIT LAKNKPILHT DLDVKSLLLE AYVKGQQELX YVVPFVAKVL
ESSIRSVVFR PPNPWTMAIM NVLAELHQEH HHHHH
```

**SUMO-GW182 SD**

$His_6$-SUMO-linker-GW182(1260-1690)

```
MGSSHHHHHH SSGLVPRGSH MASMSDSEVN QEAKPEVKPE VKPETHINLK VSDGSSEIFF
KIKKTTPLRR LMEAFAKRQG KEMDSLRFLY DGIRIQADQT PEDLDMEDND IIEAHREQIG
GSEFNTFAPY PLAGLNPNMN VNSMDMTGGL SVKDPSQSQS RLPQWTHPNS MDNLPSAASP
LEQNPSKHGA IPGGLSIGPP GKSSIDDSYG RYDLIQNSES PASPPVAVPH SWSRAKSDSD
KISNGSSINW PPEFHPGVPW KGLQNIDPEN DPDVTPGSVP TGPTINTTIQ DVNRYLLKSG
GKLSDIKSTW SSGPTSHTQA SLSHELWKVP RNSTAPTRPP PGLTNPKPSS TWGASPLGWT
SSYSSGSAWS TDTSGRTSSW LVLRNLTPQI DGSTLRTLCL QHGPLITFHL NLTQGNAVVR
YSSKEEAAKA QKSLHMCVLG NTTILAEFAG EEEVNRFLAQ GQALPPTSSW QSSSASSQPR
LSAAGSSHGL VRSDAGHWNA PCLGGKGSSE LLWGGVPQYS SSLWGPPSAD DSRVIGSPTP
LTTLLPGDLL SGESL
```

**SUMO-GW182 SD10**

$His_6$-SUMO-linker-GW182SD10(1260-1620)-FLAG

```
MGSSHHHHHH SSGLVPRGSH MASMSDSEVN QEAKPEVKPE VKPETHINLK VSDGSSEIFF
KIKKTTPLRR LMEAFAKRQG KEMDSLRFLY DGIRIQADQT PEDLDMEDND IIEAHREQIG
GSEFNTFAPY PLAGLNPNMN VNSMDMTGGL SVKDPSQSQS RLPQWTHPNS MDNLPSAASP
LEQNPSKHGA IPGGLSIGPP GKSSIDDSYG RYDLIQNSES PASPPVAVPH SWSRAKSDSD
KISNGSSINW PPEFHPGVPW KGLQNIDPEN DPDVTPGSVP TGPTINTTIQ DVNRYLLKSG
GKLSDIKSTW SSGPTSHTQA SLSHELWKVP RNSTAPTRPP PGLTNPKPSS TWGASPLGWT
SSYSSGSAWS TDTSGRTSSW LVLRNLTPQI DGSTLRTLCL QHGPLITFHL NLTQGNAVVR
YSSKEEAAKA QKSLHMCVLG NTTILAEFAG EEEVNRFLAQ GQALPPTSSW QSSSASSQPR
LSAAGADYKD DDDK
```

**GW182 SD10**

linker-GW182SD10(1260-1620)-FLAG

```
SEFNTFAPYP LAGLNPNMNV NSMDMTGGLS VKDPSQSQSR LPQWTHPNSM DNLPSAASPL
EQNPSKHGAI PGGLSIGPPG KSSIDDSYGR YDLIQNSESP ASPPVAVPHS WSRAKSDSDK
ISNGSSINWP PEFHPGVPWK GLQNIDPEND PDVTPGSVPT GPTINTTIQD VNRYLLKSGG
KLSDIKSTWS SGPTSHTQAS LSHELWKVPR NSTAPTRPPP GLTNPKPSST WGASPLGWTS
SYSSGSAWST DTSGRTSSWL VLRNLTPQID GSTLRTLCLQ HGPLITFHLN LTQGNAVVRY
SSKEEAAKAQ KSLHMCVLGN TTILAEFAGE EEVNRFLAQG QALPPTSSWQ SSSASSQPRL
SAAGADYKDD DDK
```

**PARN C-mCherry**

PARN(500-639)-linker-mCherry

```
YAESYRIQTY AEYMGRKQEE KQIKRKWTED SWKEADSKRL NPQCIPYTLQ NHYYRNNSFT
APSTVGKRNL SPSQEEAGLE DGVSGEISDT ELEQTDSCAE PLSEGRKKAK KLKRMKKELS
PAGSISKNSP ATLFEVPDTW LEVLFQGPGS AGSAAGSGEF VSKGEEDNMA IIKEFMRFKV
HMEGSVNGHE FEIEGEGEGR PYEGTQTAKL KVTKGGPLPF AWDILSPQFM YGSKAYVKHP
```

ADIPDYLKLS FPEGFKWERV MNFEDGGVVT VTQDSSLQDG EFIYKVKLRG TNFPSDGPVM
QKKTMGWEAS SERMYPEDGA LKGEIKQRLK LKDGGHYDAE VKTTYKAKKP VQLPGAYNVN
IKLDITSHNE DYTIVEQYER AEGRHSTGGM DELYK

**CNOT1 M short**

linker-CNOT1(800-999)

SEFNNDPFVQ RKLGTSGLNQ PTFQQTDLSQ VWPEANQHFS KEIDDEANSY FQRIYNHPPH
PTMSVDEVLE MLQRFKDSTI KREREVFNCM LRNLFEEYRF FPQYPDKELH ITACLFGGII
EKGLVTYMAL GLALRYVLEA LRKPFGSKMY YFGIAALDRF KNRLKDYPQY CQHLASISHF
MQFPHHLQEY IEYGQQSRDP PVK

**SUMO-GW182 SD peptide**

$His_6$-SUMO-linker-GW182(1320-1366)

MGSSHHHHHH SSGLVPRGSH MASMSDSEVN QEAKPEVKPE VKPETHINLK VSDGSSEIFF
KIKKTTPLRR LMEAFAKRQG KEMDSLRFLY DGIRIQADQT PEDLDMEDND IIEAHREQIG
GSEFPSKHGA IPGGLSIGPP GKSSIDDSYG RYDLIQNSES PASPPVAVPH S

**Supplementary Table S1.**

Data of intrinsically disordered proteins collected from literature and shown in Figure 3.

| No. | Protein | N (res) | M (Da) | $M^{1/3}$ ($Da^{1/3}$) | $R_H$ (Å) | Method | Varying conditions | Ref. |
|---|---|---|---|---|---|---|---|---|
| 1 | Aß40 | 40 | 4330 | 16.30 | 9.0 | FCS | | (Nag et al. 2011) |
| 2 | Aß42 | 42 | 4514 | 16.53 | 9.0 | FCS | | (Nag et al. 2011) |
| 3 | Smad binding domain (SBD) | 61 | 6264 | 18.43 | 25.6 | PFG-NMR | | (Chong et al. 2004) |
| 4 | EHD-L16A | 61 | 7408 | 19.49 | 20.1 | AUC | | (Mayor et al. 2003) |
| 5 | p57-ID | 71 | 8271 | 20.22 | 24.0 | SEC | | (Adkins and Lumb 2002) |
| 6 | p53-TAD | 73 | 8205 | 20.17 | 23.8 | SEC | | (Lowry et al. 2008) |
| 7 | $Ntr2^{1-75}$ | 75 | 8418 | 20.34 | 21.0 | DLS | | (Wollenhaupt et al. 2018) |
| 8 | PDE-γ | 87 | 9669 | 21.30 | 24.8 | SEC | | (Uversky et al. 2002b) |
| 9 | C-term-Vmw65 | 89 | 9330 | 21.05 | 28.0 | SEC | | (Donaldson and Capone 1992) |
| 10 | p53(1-93) | 93 | 9979 | 21.53 | 32.8 | DLS | | (Perez et al. 2014) |
| 11 | p53(1-93) $ALA^-$ | 93 | 9811 | 21.41 | 31.1 | DLS | | (Perez et al. 2014) |
| 12 | p53(1-93) $PRO^-$ | 93 | 9098 | 20.88 | 27.5 | DLS | | (Perez et al. 2014) |
| 13 | p53(1-93) $PRO^-$ $ALA^-$ | 93 | 8929 | 20.75 | 27.7 | DLS | | (Perez et al. 2014) |
| 14 | p53(1-93) | 93 | 9979 | 21.53 | 32.0 | SEC | | (Perez et al. 2014) |
| 15 | p53(1-93) $ALA^-$ | 93 | 9811 | 21.41 | 30.4 | SEC | | (Perez et al. 2014) |
| 16 | p53(1-93) $PRO^-$ | 93 | 9098 | 20.88 | 27.4 | SEC | | (Perez et al. 2014) |
| 17 | p53(1-93) $PRO^-$ $ALA^-$ | 93 | 8929 | 20.75 | 27.4 | SEC | | (Perez et al. 2014) |
| 18 | p53(1-93) | 93 | 9979 | 21.53 | 35.0 | DLS | 5 °C | (Langridge et al. 2014) |
| 19 | p53(1-93) | 93 | 9979 | 21.53 | 26.0 | DLS | 75 °C | (Langridge et al. 2014) |
| 20 | $E_m$ protein | 93 | 9963 | 21.52 | 28.2 | SEC | | (McCubbin et al. 1985) |
| 21 | Mlph(147-240) | 94 | 10111 | 21.62 | 28.0 | SEC | | (Geething and Spudich 2007) |
| 22 | Hdm2-ABD | 95 | 10503 | 21.90 | 25.7 | PFG-NMR | | (Sivakolundu et al. 2008) |
| 23 | Sm1 | 104 | 11834 | 22.79 | 23.4 | PFG-NMR | | (Danielsson et al. 2008) |
| 24 | C-terminal fragment TyrRS(Δ4) | 107 | 11908 | 22.84 | 21.0 | SEC | | (Guez et al. 2000) |
| 25 | His-Ek-LjIDP1 | 108 | 11680 | 22.69 | 24.5 | SEC | | (Haaning et al. 2008) |
| 26 | prothymosin α (1) | 109 | 11985 | 22.88 | 31.4 | SEC | pH 7.5 | (Uversky et al. 1999) |
| 27 | prothymosin α (1) | 109 | 11985 | 22.88 | 24.9 | SEC | pH 2.2 | (Uversky et al. 1999) |
| 28 | prothymosin α (2) | 110 | 12074 | 22.94 | 33.7 | PFG-NMR | no $Zn^{2+}$ | (Yi et al. 2007) |
| 29 | prothymosin α (2) | 110 | 12074 | 22.94 | 24.1 | PFG-NMR | $nZn^{2+}$/nProTα = 30 | (Yi et al. 2007) |
| 30 | ASR1 | 115 | 13130 | 23.59 | 27.4 | SEC | | (Goldgur et al. 2007) |

| | | | | | | | | |
|---|---|---|---|---|---|---|---|---|
| 31 | Nup116 FG Domain | 126 | 12613 | 23.28 | 25.2 | SEC | | (Krishnan et al. 2008) |
| 32 | Nup116 FG Domain F>A | 126 | 11852 | 22.80 | 27.1 | SEC | | (Krishnan et al. 2008) |
| 33 | TC1 | 126 | 14714 | 24.50 | 26.5 | PFG-NMR | | (Gall et al. 2007) |
| 34 | γ-synuclein | 127 | 13331 | 23.71 | 30.4 | SEC | pH 7.5, 100 mM NaCl | (Uversky et al. 2002a) |
| 35 | γ-synuclein | 127 | 13331 | 23.71 | 26.5 | SEC | pH 3.0, 100 mM NaCl | (Uversky et al. 2002a) |
| 36 | AaFEcR | 131 | 13295 | 23.69 | 27.7 | SEC | nM $Ca^{2+}$/nAaFEcR = 0.05 | (Więch et al. 2019) |
| 37 | AaFEcR | 131 | 13295 | 23.69 | 24.4 | SEC | nM $Zn^{2+}$/nAaFEcR = 2.5 | (Więch et al. 2019) |
| 38 | ß-synuclein | 134 | 14288 | 24.27 | 33.9 | SEC | pH 7.5, 100 mM NaCl | (Uversky et al. 2002a) |
| 39 | ß-synuclein | 134 | 14288 | 24.27 | 27.5 | SEC | pH 3.0, 100 mM NaCl | (Uversky et al. 2002a) |
| 40 | CaD136 | 136 | 14427 | 24.34 | 28.1 | SEC | | (Permyakov et al. 2003) |
| 41 | $N_{TAIL}$ | 139 | 15399 | 24.88 | 27.0 | SEC | | (Longhi et al. 2003) |
| 42 | $N_{TAIL}$ | 139 | 15399 | 24.88 | 30.0 | DLS | | (Longhi et al. 2003) |
| 43 | α-synuclein | 140 | 14460 | 24.36 | 28.2 | PFG-NMR | | (Paleologou et al. 2008) |
| 44 | S87A α-synuclein | 140 | 14444 | 24.35 | 28.1 | PFG-NMR | | (Paleologou et al. 2008) |
| 45 | P Ser-129 Ser-87 α-synuclein | 140 | 14616 | 24.45 | 35.3 | PFG-NMR | | (Paleologou et al. 2008) |
| 46 | P Ser-129 S87A α-synuclein | 140 | 14522 | 24.40 | 34.7 | PFG-NMR | | (Paleologou et al. 2008) |
| 47 | α-synuclein | 140 | 14460 | 24.36 | 31.8 | SEC | pH 7.5, 100 mM NaCl | (Uversky et al. 2002a) |
| 48 | α-synuclein | 140 | 14460 | 24.36 | 27.9 | SEC | pH 3.0, 100 mM NaCl | (Uversky et al. 2002a) |
| 49 | hNL3-cyt | 140 | 15290 | 24.82 | 28.3 | SEC | | (Paz et al. 2008) |
| 50 | hNL3-cyt | 140 | 15290 | 24.82 | 23.3 | DLS | | (Paz et al. 2008) |
| 51 | hNL3-cyt | 140 | 15290 | 24.82 | 25.0 | FCS | | (Paz et al. 2008) |
| 52 | hNL3-cyt | 140 | 15290 | 24.82 | 27.3 | AUC | | (Paz et al. 2008) |
| 53 | ShB-C | 146 | 15912 | 25.15 | 32.9 | SEC | | (Magidovich et al. 2007) |
| 54 | $Ntr2^{1\text{-}162}$ | 162 | 18141 | 26.28 | 31.0 | DLS | | (Wollenhaupt et al. 2018) |
| 55 | Fos-AD | 168 | 17612 | 26.02 | 35.0 | SEC | | (Campbell et al. 2000) |
| 56 | HIF-1α (530-698) | 169 | 18645 | 26.52 | 38.3 | SEC | | (Sánchez-Puig et al. 2005a) |
| 57 | CFTR R region | 189 | 21410 | 27.77 | 32.0 | PFG-NMR | | (Baker 2009) |
| 58 | Tau K32 | 198 | 21030 | 27.60 | 45.0 | PFG-NMR | | (Soragni et al. 2008) |
| 59 | HIF-1α (403-603) | 201 | 22157 | 28.09 | 44.3 | SEC | | (Sánchez-Puig et al. 2005a) |
| 60 | DARPP-32 | 202 | 22614 | 28.28 | 34.0 | SEC | | (Hemmings et al. 1984) |
| 61 | Securin | 204 | 22247 | 28.12 | 39.7 | SEC | | (Sánchez-Puig et al. 2005b) |
| 62 | SNAP25 | 206 | 23315 | 28.57 | 39.3 | DLS | | (Choi et al. 2011) |
| 63 | Gliotactin-cyt | 217 | 23594 | 28.68 | 33.4 | SEC | | (Zeev-Ben-Mordehai et al. 2003) |

| | | | | | | | | |
|---|---|---|---|---|---|---|---|---|
| 64 | 3D7-6H MSP2 | 237 | 24191 | 28.92 | 41.0 | PFG-NMR | pH 3.6; 10 mM HOAc | (Zhang et al. 2008) |
| 65 | 3D7-6H MSP2 | 237 | 24191 | 28.92 | 34.3 | PFG-NMR | pH 7.0; PBS | (Zhang et al. 2008) |
| 66 | 3D7-6H MSP2 | 237 | 24191 | 28.92 | 34.9 | PFG-NMR | pH 3.5; 10 mM HOAc + 135 mM NaCl | (Zhang et al. 2008) |
| 67 | RYBP | 234 | 25644 | 29.49 | 39.5 | PFG-NMR | | (Neira et al. 2009) |
| 68 | $H_6$-PNT | 236 | 24849 | 29.18 | 41.0 | SEC | | (Karlin et al. 2002) |
| 69 | $H_6$-PNT | 236 | 24849 | 29.18 | 47.0 | DLS | | (Karlin et al. 2002) |
| 70 | Mlph(147-403) | 257 | 27573 | 30.21 | 46.0 | DLS | | (Geething and Spudich 2007) |
| 71 | Mlph(147-403) | 257 | 27573 | 30.21 | 49.0 | SEC | | (Geething and Spudich 2007) |
| 72 | $Ntr2^{FL}$ | 322 | 36647 | 33.22 | 42.0 | DLS | | (Wollenhaupt et al. 2018) |
| 73 | DBE | 345 | 39843 | 34.15 | 33.1 | SEC | | (Yiu et al. 2006) |
| 74 | Calreticulin | 404 | 46791 | 36.03 | 46.2 | SEC | | (Bouvier and Stafford 2000) |
| 75 | HeV PNT | 410 | 45216 | 35.63 | 44.0 | SEC | | (Habchi et al. 2010) |
| 76 | HeV PNT | 410 | 45216 | 35.63 | 50.0 | DLS | | (Habchi et al. 2010) |
| 77 | NiV PNT | 412 | 45331 | 35.66 | 44.0 | SEC | | (Habchi et al. 2010) |
| 78 | NiV PNT | 412 | 45331 | 35.66 | 44.0 | DLS | | (Habchi et al. 2010) |
| 79 | Nup159pΔNΔC | 441 | 44698 | 35.49 | 61.0 | SEC | | (Denning et al. 2003) |
| 80 | Mid1p-N452 | 458 | 50503 | 36.96 | 49.0 | SEC | | (Chatterjee and Pollard 2019) |
| 81 | Starmaker | 593 | 64043 | 40.01 | 78.6 | SEC | | (Kapłon et al. 2008) |
| 82 | Nsp1pΔC | 603 | 61887 | 39.55 | 74.0 | SEC | | (Denning et al. 2003) |
| 83 | OMM-64 | 608 | 64473 | 40.10 | 76.2 | SEC | | (Poznar et al. 2017) |
| 84 | OMM-64 | 608 | 64473 | 40.10 | 74.5 | FCS | | (Poznar et al. 2017) |
| 85 | OMM-64 | 608 | 64473 | 40.10 | 75.9 | AUC | | (Poznar et al. 2017) |
| 86 | OMM-64 | 608 | 64473 | 40.10 | 85.0 | SEC | no salt | (Poznar et al. 2017) |
| 87 | OMM-64 | 608 | 64473 | 40.10 | 70.0 | SEC | 100 mM $CaCl_2$ | (Poznar et al. 2017) |
| 88 | Nup100pΔC | 640 | 64454 | 40.09 | 59.0 | SEC | | (Denning et al. 2003) |
| 89 | Nup2p | 720 | 77881 | 42.70 | 79.0 | SEC | | (Denning et al. 2002) |
| 90 | Nup85p | 744 | 84898 | 43.95 | 51.0 | SEC | | (Denning et al. 2003) |
| 91 | Caldesmon | 771 | 88747 | 44.61 | 91.0 | SEC | | (Lynch et al. 1987) |
| 92 | Nup1pΔN | 777 | 80671 | 43.21 | 75.0 | SEC | | (Denning et al. 2003) |
| 93 | Fesselin | 996 | 107712 | 47.58 | 53.0 | SEC | | (Khaymina et al. 2007) |

### Sequences of proteins gathered in Table S1.

**Aβ40**
DAEFRHDSGY EVHHQKLVFF AEDVGSNKGA IIGLMVGGVV

**Aβ42**
DAEFRHDSGY EVHHQKLVFF AEDVGSNKGA IIGLMVGGVV IA

**Smad binding domain (SBD)**
GSMMSASSQS PNPNNPAEYC STIPPLEYCS TIPPLQQAQA SGALSSPPPT VMVPVGVLKH P

**EHD-L16A**
TNDEKRPRTA FSSEQAARLK REFNENRYLT ERRRQQLSSE LGLNEAQIKI WFQNKRAKIK K

**p57-ID**
TSACRSLFGP VDHEELSREL QARLAELNAE DQNRWDYDFQ QDMPLRGPGR LQWTEVDSDS
VPAFYRETVQ V

**p53-TAD**
MEEPQSDPSV EPPLSQETFS DLWKLLPENN VLSPLPSQAM DDLMLSPDDI EQWFTEDPGP
DEAPRMPEAA PRV

**Ntr2$^{1-75}$**
MAIKKRNKIR LPSGSPEEVG IDGSAHKPMQ QIKPLVSNDS EDDDNDICVL QPIKFKKVPK
RDITFDGEQA IKEDN

**PDE-γ**
MNLEPPKAEI RSATRVMGGP VTPRKGPPKF KQRQTRQFKS KPPKKGVQGF GDDIPGMEGL
GTDITVICPW EAFNHLELHE LAQYGII

**C-term-Vmw65**
GSAGHTRRLS TAPPTDVSLG DELHLDGEDV AMAHADALDD FDLDMLGDGD SPGPGFTPHD
SAPYGALDMA DFEFEQMFTD ALGIDEYGG

**p53(1-93)**
MEEPQSDPSV EPPLSQETFS DLWKLLPENN VLSPLPSQAM DDLMLSPDDI EQWFTEDPGP
DEAPRMPEAA PPVAPAPAAP TPAAPAPAPS WPL

**p53(1-93) ALA-**
MEEPQSDPSV EPPLSQETFS DLWKLLPENN VLSPLPSQGM DDLMLSPDDI EQWFTEDPGP
DEGPRMPEGG PPVGPGPGGP TPGGPGPGPS WPL

**p53(1-93) PRO-**
MEEGQSDGSV EGGLSQETFS DLWKLLGENN VLSGLGSQAM DDLMLSGDDI EQWFTEDGGG
DEAGRMGEAA GGVAGAGAAG TGAAGAGAGS WGL

**p53(1-93) ALA- PRO-**
MEEGQSDGSV EGGLSQETFS DLWKLLGENN VLSGLGSQGM DDLMLSGDDI EQWFTEDGGG
DEGGRMGEGG GGVGGGGGGG TGGGGGGGGS WGL

**$E_m$ protein**
MASGQQERSQ LDRKAREGET VVPGGTGGKS LEAQENLAEG RSRGGQTRRE QMGEEGYSQM
GRKGGLSTND ESGGDRAARE GIDIDESKFK TKS

**Mlph(147-240)**
GGGGSEPSLE EGNGDSEQTD EDGDLDTEAR DQPLNSKKKK RLLSFRDVDF EEDSDHLVQP
CSQTLGLSSV PESAHSLQSL SGEPYSEDTT SLEP

**Hdm2-ABD**
SSSSESTGTP SNPDLDAGVS EHSGDWLDQD SVSDQFSVEF EVESLDSEDY SLSEEGQELS
DEDDEVYQVT VYQAGESDTD SFEEDPEISL ADYWK

**Sm1**
MQNSQDYFYA QNRCQQQQAP STLRTVTMAE FRRVPLPPMA EVPMLSTQNS MGSSASASAS
SLEMWEKDLE ERLNSIDHDM NNNKFGSGEL KSMFNQGKVE EMDF

**C-terminal fragment TyrRS(Δ4)**
ALFSGDIANL TAAEIEQGFK DVPSFVHEGG DVPLVELLVS AGISPSKRQA REDIQNGAIY
VNGERLQDVG AILTAEHRLE GRFTVIRRGK KKYYLIRYAL GHHHHHH

**His-Ek-LjIDP1**
MAHHHHHHVD DDDKMARSFT NIKAISALVA EEFSNSLARR GYAATAQSAG RVGASMSGKM
GSTKSGEEKA AAREKVSWVP DPVTGYYKPE NIKEIDVAEL RSAVLGKN

**prothymosin α (1)**
SDAAVDTSSE ITTKDLKEKK EVVEEAENGR DAPANGNANE ENGEQEADNE VDEEEEEGGE
EEEEEEEGDG EEEDVDEDEE AESATGKRAA EDDEDDDVDT KKQKTDEDD

**prothymosin α (2)**
MSDAAVDTSS EITTKDLKEK KEVVEEAENG RDAPANGNAN EENGEQEADN EVDEEEEEGG
EEEEEEEEGD GEEEDGDEDE EAESATGKRA AEDDEDDDVD TKKQKTDEDD

**ASR1**
MEEEKHHHHH LFHHKDKAEE GPVDYEKEIK HHKHLEQIGK LGTVAAGAYA LHEKHEAKKD
PEHAHKHKIE EEIAAAAAVG AGGFAFHEHH EKKDAKKEEK KKLRGDTTIS SKLLF

**Nup116 FG Domain**
GSRRASVGSG ALFGAKPASG GLFGQSAGSK AFGMNTNPTG TTGGLFGQTN QQQSGGGLFG
QQQNSNAGGL FGQNNQSQNQ SGLFGQQNSS NAFGQPQQQG GLFGSKPAGG LFGQQQGAST
HHHHHH

**Nup116 FG Domain F>A**
GSRRASVGSG ALAGAKPASG GLAGQSAGSK AAGMNTNPTG TTGGLAGQTN QQQSGGGLAG
QQQNSNAGGL AGQNNQSQNQ SGLAGQQNSS NAAGQPQQQG GLAGSKPAGG LAGQQQGAST
HHHHHH

**TC1**
HHHHHHXXXX XXXXXXXXXX MKAKRSHQAI IMSTSLRVSP SIHGYHFDTA SRKKAVGNIF
ENTDQESLER LFRNSGDKKA EERAKIIFAI DQDVEEKTRA LMALKKRTKD KLFQFLKLRK
YSIKVH

**γ-synuclein**
MDVFKKGFSI AKEGVVGAVE KTKQGVTEAA EKTKEGVMYV GAKTKENVVQ SVTSVAEKTK
EQANAVSEAV VSSVNTVATK TVEEAENIAV TSGVVRKEDL RPSAPQQEGE ASKEKEEVAE
EAQSGGD

**AaFEcR**
GPSAGLVPRG SGGIEGRHML EEIWDVQDIP PSMQAQMHSH GTQSSSSSSS SSSSSSNGSS
NGNSSSNSNS SQHGPHPHPH GQQLTPNQQQ HQQQHSQLQQ VHANGSGSGG GSNNNSSSGG
VVPGLGMLDQ V

**β-synuclein**
MDVFMKGLSM AKEGVVAAAE KTKQGVTEAA EKTKEGVLYV GSKTREGVVQ GVASVAEKTK
EQASHLGGAV FSGAGNIAAA TGLVKREEFP TDLKPEEVAQ EAAEEPLIEP LMEPEGESYE
DPPQEEYQEY EPEA

**CaD136**
RLEQYTSAVV GNKAAKPAKP AASDLPVPAE GVRNIKSMWE KGNVFSSPGG TGTPNKETAG
LKVGVSSRIN EWLTKTPEGN KSPAPKPSDL RPGDVSGKRN LWEKQSVEKP AASSSKVTAT

GKKSETNGLR QFEKEP

**N$_{TAIL}$**
MRGSHHHHHH XXXHTTEDKI SRAVGPRQAQ VSFLHGDQSE NELPRLGGKE DRRVKQSRGE
ARESYRETGP SRASDARAAH LPTGTPLDID TASESSQDPQ DSRRSADALL RLQAMAGISE
EQGSDTDTPI VYNDRNLLD

**α-synuclein**
MDVFMKGLSK AKEGVVAAAE KTKQGVAEAA GKTKEGVLYV GSKTKEGVVH GVATVAEKTK
EQVTNVGGAV VTGVTAVAQK TVEGAGSIAA ATGFVKKDQL GKNEEGAPQE GILEDMPVDP
DNEAYEMPSE EGYQDYEPEA

**S87A α-synuclein**
MDVFMKGLSK AKEGVVAAAE KTKQGVAEAA GKTKEGVLYV GSKTKEGVVH GVATVAEKTK
EQVTNVGGAV VTGVTAVAQK TVEGAGAIAA ATGFVKKDQL GKNEEGAPQE GILEDMPVDP
DNEAYEMPSE EGYQDYEPEA

**hNL3-cyt**
MGSSHHHHHH SSGLVPRGSH MAYRKDKRRQ EPLRQPSPQR GAGAPELGAA PEEELAALQL
GPTHHECEAG PPHDTLRLTA LPDYTLTLRR SPDDIPLMTP NTITMIPNSL VGLQTLHPYN
TFAAGFNSTG LPHSHSTTRV

**ShB-C**
MXXGQHMKKS SLSESSSDMM DLDDGVESTP GLTETHPGRS AVAPFLGAQQ QQQQPVASSL
SMSIDKQLQH PLQQLTQTQL YQQQQQQQQQ QQNGFKQQQQ QTQQQLQQQQ SHTINASAAA
ATSGSGSSGL TMRHNNALAV SIETDV

**Ntr2$^{1\text{-}162}$**
MAIKKRNKIR LPSGSPEEVG IDGSAHKPMQ QIKPLVSNDS EDDDNDICVL QPIKFKKVPK
RDITFDGEQA IKEDNSHYED LYHSKKNTNA STRNKDDLLI LNMEDLMEGN HHLLSDSSEA
GSSSEGEHIS SIPTRGEIAK LKAQKSLSRR KISESDVTTE RD

**Fos-AD**
GSHMSVASLD LTGGLPEVAT PESEEAFTLP LLNDPEPKPS VEPVKSISSM ELKTEPFDDF
LFPASSRPSG SETARSVPDM DLSGSFYAAD WEPLHSGSLG MGPMATELEP LCTPVVTCTP
SCTAYTSSFV FTYPEADSFP SCAAAHRKGS SSNEPSSDSL SSPTLLAL

**HIF-1α (530-698)**
EFKLELVEKL FAEDTEAKNP FSTQDTDLDL EMLAPYIPMD DDFQLRSFDQ LSPLESSSAS
PESASPQSTV TVFQQTQIQE PTANATTTTA TTDELKTVTK DRMEDIKILI ASPSPTHIHK
ETTSATSSPY RDTQSRTASP NRAGKGVIEQ TEKSHPRSPN VLSVALSQR

**CFTR R region**
GAMESAERRN SILTETLHRF SLEGDAPVSW TETKKQSFKQ TGEFGEKRKN SILNPINSIR
KFSIVQKTPL QMNGIEEDSD EPLERRLSLV PDSEQGEAIL PRISVISTGP TLQARRRQSV
LNLMTHSVNQ GQNIHRKTTA STRKVSLAPQ ANLTELDIYS RRLSQETGLE ISEEINEEDL
KECLFDDME

**Tau K32**
MSSPGSPGTP GSRSRTPSLP TPPTREPKKV AVVRTPPKSP SSAKSRLQTA PVPMPDLKNV
KSKIGSTENL KHQPGGGKVQ IINKKLDLSN VQSKCGSKDN IKHVPGGGSV QIVYKPVDLS
KVTSKCGSLG NIHHKPGGGQ VEVKSEKLDF KDRVQSKIGS LDNITHVPGG GNKKIETHKL
TFRENAKAKT DHGAEIVY

**HIF-1α (403-603)**
AAGDTIISLD FGSNDTETDD QQLEEVPLYN DVMLPSPNEK LQNINLAMSP LPTAETPKPL
RSSADPALNQ EVALKLEPNP ESLELSFTMP QIQDQTPSPS DGSTRQSSPE PNSPSEYCFY
VDSDMVNEFK LELVEKLFAE DTEAKNPFST QDTDLDLEML APYIPMDDDF QLRSFDQLSP
LESSSASPES ASPQSTVTVF Q

**DARPP-32**
MDPKDRKKIQ FSVPAPPSQL DPRQVEMIRR RRPTPAMLFR LSEHSSPEEE ASPHQRASGE
GHHLKSKRSN PCAYTPPSLK AVQRIAESHL QSISNLGENQ ASEEEDELGE LRELGYPREE
EEEEEEEDEE EEEDSQAEVL KGSRGSAGQK TTYGQGLEGP WERPPPLDGP QRDGSSEDQV
EDPALNEPGE EPQRPAHPEP GT

**Securin**
XXMATLIYVD KENGEPGTRV VAKDGLKLGS GPSIKALDGR SQVSTPRFGK TFDAPPALPK
ATRKALGTVN RATEKSVKTK GPLKQKQPSF SAKKMTEKTV KAKSSVPASD DAYPEIEKFF
PFNPLDFESF DLPEEHQIAH LPLSGVPLMI LDEERELEKL FQLGPPSPVK MPSPPWESNL
LQSPSSILST LDVELPPVCC DIDI

**SNAP25**
MAEDADMRNE LEEMQRRADQ LADESLESTR RMLQLVEESK DAGIRTLVML DEQGEQLERI
EEGMDQINKD MKEAEKNLTD LGKFCGLCVC PCNKLKSSDA YKKAWGNNQD GVVASQPARV
VDEREQMAIS GGFIRRVTND ARENEMDENL EQVSGIIGNL RHMALDMGNE IDTQNRQIDR
IMEKADSNKT RIDEANQRAT KMLGSG

**Gliotactin-cyt**
XXXRNAKRQS DRFYDEDVFI NGEGLEPEQD TRGVDNAHMV TNHHALRSRD NIYEYRDSPS
TKTLASKAHT DTTSLRSPSS LAMTQKSSSQ ASLKSGISLK ETNGHLVKQS ERAATPRSQQ
NGSIAKVASP PVEEKRLLQP LSSTPVTQLQ AEPAKRVPTA ASVSGSSRST TPVPSARSTT
THTTTATLSS QPAAQPRRTH LVEGVPQTSV XHHHHHH

**3D7-6H MSP2**
MIKNESKYSN TFINNAYNMS IRRSMAESKP STGAGGSAGG SAGGSAGGSA GGSAGGSAGS
GDGNGADAEG SSSTPATTTT TKTTTTTTTT NDAEASTSTS SENPNHKNAE TNPKGKGEVQ
EPNQANKETQ NNSNVQQDSQ TKSNVPPTQD ADTKSPTAQP EQAENSAPTA EQTESPELQS
APENKGTGQH GHMHGSRNNH PQNTSDSQKE CTDGNKENCG AATSLLNNSS NHHHHHH

**RYBP**
HHHHHHMTMG DKKSPTRPKR QAKPAADEGF WDCSVCTFRN SAEAFKCSIC DVRKGTSTRK
PRINSQLVAQ QVAQQYATPP PPKKEKKEKV EKQDKEKPEK DKEISPSVTK KNTNKKTKPK
SDILKDPPSE ANSIQSANAT TKTSETNHTS RPRLKNVDRS TAQQLAVTVG NVTVIITDFK
EKTRSSSTSS STVTSSAGSE QQNQSSSGSE STDKGSSRSS TPKGDMSAVN DESF

**$H_6$-PNT**
HHHHHHMAEE QARHVKNGLE CIRALKAEPI GSLAIEEAMA AWSEISDNPG QERATCREEK
AGSSGLSKPC LSAIGSTEGG APRIRGQGPG ESDDDAETLG IPPRNLQASS TGLQCYYVYD
HSGEAVKGIQ DADSIMVQSG LDGDSTLSGG DNESENSDVD IGEPDTEGYA ITDRGSAPIS
MGFRASDVET AEGGEIHELL RLQSRGNNFP KLGKTLNVPP PPDPGRASTS GTPIKK

**Mlph(147-403)**
GGGGSEPSLE EGNGDSEQTD EDGDLDTEAR DQPLNSKKKK RLLSFRDVDF EEDSDHLVQP
CSQTLGLSSV PESAHSLQSL SGEPYSEDTT SLEPEGLEET GARALGCRPS PEVQPCSPLP
SGEDAHAELD SPAASCKSAF GTTAMPGTDD VRGKHLPSQY LADVDTSDED SIQGPRAASQ
HSKRRARTVP ETQILELNKR MSAVEHLLVH LENTVLPPSA QEPTVETHPS ADTEEETLRR
RLEELTSNIS GSSTSSE

**$Ntr2^{FL}$**
MAIKKRNKIR LPSGSPEEVG IDGSAHKPMQ QIKPLVSNDS EDDDNDICVL QPIKFKKVPK
RDITFDGEQA IKEDNSHYED LYHSKKNTNA STRNKDDLLI LNMEDLMEGN HHLLSDSSEA
GSSSEGEHIS SIPTRGEIAK LKAQKSLSRR KISESDVTTE RDYVKLLDSE DKREIMETIR
LNGGLKRNNE KEITNFSDDE MQGFQDEMLA LTDNQIAIQK DSKRKIIEKA INEVPYRTNE
EWETQLLSKG NINKSNEKII TPLPVLFPDD DESGNSIERI NEMVSKICLQ RKKVEMRLQA
LEKTKIDLEK SKASLINKLI GN

**DBE**
MSESEAEETK ISTEPVDNAW SMKIPAFRQE DNPHGMVEES SFATLFPKYR ERYLKEVWPL
VEQCLAEHHL KAELDLMEGS MVVKTSRKTW DPYIIIKARD MIKLMARSVP FEQAKRVLQD
DIGCDIIKIG NLVHKKEKFV KRRQRLIGPN GATLKSIELL TDCYVLVQGN TVSALGPYKG

LQQVRDIVLE TMNNVHPIYN IKALMIKREL MKDPRLANED WSRFLPKFKN KNISKRKQPK
VKKQKKEYTP FPPSQPESKV DKQLASGEYF LNQEQKQAKR NQERTEKQKE AAKRQDERRN
KDFVPPTEES AASSRKKEDG SSSSKVDVKA LKAKLIKANK KARSS

**Calreticulin**
GIPGEPAVYF KEQFLDGDGW TSRWIESKHK SDFGKFVLSS GKFYGDEEKD KGLQTSQDAR
FYALSASFEP FSNKGQTLVV QFTVKHEQNI DCGGGYVKLF PNSLDQTDMH GDSEYNIMFG
PDICGPGTKK VHVIFNYKGK NVLINKDIRC KDDEFTHLYT LIVRPDNTYE VKIDNSQVES
GSLEDDWDFL PPKKIKDPDA SKPEDWDERA KIDDPTDSKP EDWDKPEHIP DPDAKKPEDW
DEEMDGEWEP PVIQNPEYKG EWKPRQIDNP DYKGTWIHPE IDNPEYSPDP SIYAYDNFGV
LGLDLWQVKS GTIFDNFLIT NDEAYAEEFG NETWGVTKAA EKQMKDKQDE EQRLKEEEED
KKRKEEEEAE DKEDDEDKDE DEEDEEDKEE DEEEDVPGQA KDEL

**HeV PNT**
MDKLDLVNDG LDIIDFIQKN QKEIQKTYGR SSIQQPSTKD RTRAWEDFLQ STSGEHEQAE
GGMPKNDGGT EGRNVEDLSS VTSSDGTIGQ RVSNTRAWAE DPDDIQLDPM VTDVVYHDHG
GECTGHGPSS SPERGWSYHM SGTHDGNVRA VPDTKVLPNA PKTTVPEEVR EIDLIGLEDK
FASAGLNPAA VPFVPKNQST PTEEPPVIPE YYYGSGRRGD LSKSPPRGNV NLDSIKIYTS
DDEDENQLEY EDEFAKSSSE VVIDTTPEDN DSINQEEVVG DPSDQGLEHP FPLGKFPEKE
ETPDVRRKDS LMQDSCKRGG VPKRLPMLSE EFECSGSDDP IIQELEREGS HPGGSLRLRE
PPQSSGNSRN QPDRQLKTGD AASPGGVQRP GTPMPKSRIM PIKKHHHHHH

**NiV PNT**
MDKLELVNDG LNIIDFIQKN QKEIQKTYGR SSIQQPSIKD QTKAWEDFLQ CTSGESEQVE
GGMSKDDGDV ERRNLEDLSS TSPTDGTIGK RVSNTRDWAE GSDDIQLDPV VTDVVYHDHG
GECTGYGFTS SPERGWSDYT SGANNGNVCL VSDAKMLSYA PEIAVSKEDR ETDLVHLENK
LSTTGLNPTA VPFTLRNLSD PAKDSPVIAE HYYGLGVKEQ NVGPQTSRNV NLDSIKLYTS
DDEEADQLEF EDEFAGSSSE VIVGISPEDE EPSSVGGKPN ESIGRTIEGQ SIRDNLQAKD
NKSTDVPGAG PKDSAVKEEP PQKRLPMLAE EFECSGSEDP IIRELLKENS LINCQQGKDA
QPPYHWSIER SISPDKTEIV NGAVQTADRQ RPGTPMPKSR GIPIKKHHHHHH

**Nup159pΔNΔC**
SGFTFLKTQP AAANSLQSQS SSTFGAPSFG SSAFKIDLPS VSSTSTGVAS SEQDATDPAS
AKPVFGKPAF GAIAKEPSTS EYAFGKPSFG APSFGSGKSS VESPASGSAF GKPSFGTPSF
GSGNSSVEPP ASGSAFGKPS FGTPSFGSGN SSAEPPASGS AFGKPSFGTS AFGTASSNET
NSGSIFGKAA FGSSSFAPAN NELFGSNFTI SKPTVDSPKE VDSTSPFPSS GDQSEDESKS
DVDSSSTPFG TKPNTSTKPK TNAFDFGSSS FGSGFSKALE SVGSDTTFKF GTQASPFSSQ
LGNKSPFSSF TKDDTENGSL SKGSTSEIND DNEEHESNGP NVSGNDLTDS TVEQTSSTRL
PETPSDEDGE VVEEEAQKSP IGKLTETIKK SANIDMAGLK NPVFGNHVKA KSESPFSAFA
TNITKPSSTT PAFSFGNSTM N

**Mid1p-N452**
HHHHHHMKEQ EFSYREAKDV SLDSKGLENS FLSSPNREKT PLFFEGNSNE TSGYDQTKNF
THGDGDMSLG NLSELNVATD LLESLDLRSM YMHGYGHLDS SFSSQHSPDN RKRMSSTSVF
KRINSEEEGR IPSLTYSAGT MNSTSSSTAS LKGADIVADY ETFNPDQNLA ELSFDRSKSS
RKRAVEVAEF SRAKTMSPLE YTVQHPYQSH NELSTNPARA RAGSVPNLAR IPSDVKPVPP
AHLSASSTVG PRILPSLPKD TTEDNPALER VETTASLDMD YKPLEPLAPI QEAPVEDTSE
PFSSVPEATL DDSDISTESL RKKVLAKMEA KRISSGSSYA STLRKVYDFS ELSLPTNGKD
YDELYLQSSR NSEPEISTII NDSLQQENMD EDISATSIPK SQAAYGHGSV TYHEVPRYNL
TSASVGYSIS SQRGRIKSSS TIDNLSAILS SEDLRHPS

**Starmaker**
APVSNNNGTD NDESAADQRH IFTVQFNVGT PAPADGDSVT TDGKDSAEKN EAPGDSSDTT
EKPGTTDGKD SAEQHGVTTD GKDEAEQHGV TTDGQDSAEK RGEADGAPDK PDTQNGTDDT
DSDQETDASH HKTGDSDENK DKPSAEDHTD GNHAGKDSTD SKESPDTTDK PEGPDSDSAP
DGDSASAEKT DSDHSPDEDA NKSSTEADKD DTSDKDSSQT DEKHDSDASD KDEKHEDKDE
KSDEKDSSKD SEDKSQEKSD KSDDGSNSEA DEQKESVESK DHDSDSQDSD SAEKKEKHDD
KDQDSSDSAD SKDSDEDKDK DHSEQKDSED HEHKEKHTKD KEEHKDSDEG KDDEDKSKSD
EHDKDESESK EASKSDESEQ EEKKDDKSDS DNSSRDSHSD SDSDSHSDSD SDSHSDSHSD
SDSDSHSDSD SDSDSDSDSD SDSDSDSNSR DKDEKKDKSS ESRDEDSSDS DSKSNSESSE
TAEEDTNDDK DSSVEKDKTD SSDSASVEAN DSDDEHDDDS KDATPSSEDH TAEKTDEDSH

DVSDDDDDID AHDDEAGVEH GTDEASKPHQ EPDHHDDTTH GSDDGRKTSM PIS

**Nsp1pΔC**
MNFNTPQQNK TPFSFGTANN NSNTTNQNSS TGAGAFGTGQ STFGFNNSAP NNTNNANSSI
TPAFGSNNTG NTAFGNSNPT SNVFGSNNST TNTFGSNSAG TSLFGSSSAQ QTKSNGTAGG
NTFGSSSLFN NSTNSNTTKP AFGGLNFGGG NNTTPSSTGN ANTSNNLFGA TANANKPAFS
FGATTNDDKK TEPDKPAFSF NSSVGNKTDA QAPTTGFSFG SQLGGNKTVN EAAKPSLSFG
SGSAGANPAG ASQPEPTTNE PAKPALSFGT ATSDNKTTNT TPSFSFGAKS DENKAGATSK
PAFSFGAKPE EKKDDNSSKP AFSFGAKSNE DKQDGTAKPA FSFGAKPAEK NNNETSKPAF
SFGAKSDEKK DGDASKPAFS FGAKPDENKA SATSKPAFSF GAKPEEKKDD NSSKPAFSFG
AKSNEDKQDG TAKPAFSFGA KPAEKNNNET SKPAFSFGAK SDEKKDGDAS KPAFSFGAKS
DEKKDSDSSK PAFSFGTKSN EKKDSGSSKP AFSFGAKPDE KKNDEVSKPA FSFGAKANEK
KESDESKSAF SFGSKPTGKE EGDGAKAAIS FGAKPEEQKS SDTSKPAFTF GAQKDNEKKT
EES

**OMM-64**
APVNDGTEAD NDERAASLLV HLKGDKDGGG LTGSPDGVSA GTTDGTDSSK ELAGGAVDSS
PDTTDTPDAS SSDIFPDTNN RDTSVETTGN PDDSDAPDAA ESAGSQDTTD AADASEAVAE
TVDTYDIPDT DGADDREKVS TEVSTEDLDS AGVDKSPESD STESPGSDSA ESPGSDSAES
PGSDSTESPG SDSTESPRSD STDEVLTDVQ ADSADVTSDD MDEATETDKD DDKSDDKSDA
DAATDKDDSD EDKDTELDGK AHAEDTQTEE AADSDSKQGA ADSDSDTDDD RPEKDVKNDS
DDSKDTTEDD KPDKDDKKNR DSADNSNDDS DEMIQVPREE LEQQEINLKE GGVIGSQEET
VASDMEEGSD VGDQKPGPED SIEEGSPVGR QDFKHPQDSE EEELEKEAKK EKELEEAEEE
RTLKTIESDS QEDSVDESEA EPDSNSKKDI GTSDAPEPQE DDSEEDTDDS MMKEPKDSDD
AESDKDDKDK NDMDKEDMDK DDMDKDDMDK DDMDKDDVDK DASDSVDDQS ESDAEPGADS
HTVVDEIDGE ETMTPDSEEI MKSGEMDSVV EATEVPADIL DQPDQQDDMT QGASQAADAA
ATALAAQS

**Nup100pΔC**
MFGNNRPMFG GSNLSFGSNT SSFGGQQSQQ PNSLFGNSNN NNNSTSNNAQ SGFGGFTSAA
GSNSNSLFGN NNTQNNGAFG QSMGATQNSP FGSLNSSNAS NGNTFGGSSS MGSFGGNTNN
AFNNNSNSTN SPFGFNKPNT GGTLFGSQNN NSAGTSSLFG GQSTSTTGTF GNTGSSFGTG
LNGNGSNIFG AGNNSQSNTT GSLFGNQQSS AFGTNNQQGS LFGQQSQNTN NAFGNQNQLG
GSSFGSKPVG SGSLFGQSNN TLGNTTNNRN GLFGQMNSSN QGSSNSGLFG QNSMNSSTQG
VFGQNNNQMQ INGNNNNSLF GKANTFSNSA SGGLFGQNNQ QQGSGLFGQN SQTSGSSGLF
GQNNQKQPNT FTQSNTGIGL FGQNNNQQQQ STGLFGAKPA GTTGSLFGGN SSTQPNSLFG
TTNVPTSNTQ SQQGNSLFGA TKLTNMPFGG NPTANQSGSG NSLFGTKPAS TTGSLFGNNT
ASTTVPSTNG LFGNNANNST STTNTGLFGA KPDSQSKPAL GGGLFGNSNS NSSTIGQNKP
VFGGTTQNTG LFGATGTNSS AVGSTGKLFG QNNNTLNVGT QNVPPVNNTT QNALLGTTAV
PSLQQAPVTN EQLFSKISIP NSITNPVKAT TSKVNADMKR

**Nup2p**
MAKRVADAQI QRETYDSNES DDDVTPSTKV ASSAVMNRRK IAMPKRRMAF KPFGSAKSDE
TKQASSFSFL NRADGTGEAQ VDNSPTTESN SRLKALNLQF KAKVDDLVLG KPLADLRPLF
TRYELYIKNI LEAPVKSIEN PTQTKGNDAK PAKVEDVQKS SDSSSEDEVK VEGPKFTIDA
KPPISDSVFS FGPKKENRKK DESDSENDIE IKGPEFKFSG TVSSDVFKLN PSTDKNEKKT
ETNAKPFSFS SATSTTEQTK SKNPLSLTEA TKTNVDNNSK AEASFTFGTK HAADSQNNKP
SFVFGQAAAK PSLEKSSFTF GSTTIEKKND ENSTSNSKPE KSSDSNDSNP SFSFSIPSKN
TPDASKPSFS FGVPNSSKNE TSKPVFSFGA ATPSAKEASQ EDDNNNVEKP SSKPAFNLIS
NAGTEKEKES KKDSKPAFSF GISNGSESKD SDKPSLPSAV DGENDKKEAT KPAFSFGINT
NTTKTADTKA PTFTFGSSAL ADNKEDVKKP FSFGTSQPNN TPSFSFGKTT ANLPANSSTS
PAPSIPSTGF KFSLPFEQKG SQTTTNDSKE ESTTEATGNE SQDATKVDAT PEESKPINLQ
NGEEDEVALF SQKAKLMTFN AETKSYDSRG VGEMKLLKKK DDPSKVRLLC RSDGMGNVLL
NATVVDSFKY EPLAPGNDNL IKAPTVAADG KLVTYIVKFK QKEEGRSFTK AIEDAKKEMK

**Nup85p**
MTIDDSNRLL MDVDQFDFLD DGTAQLSNNK TDEEEQLYKR DPVSGAILVP MTVNDQPIEK
NGDKMPLKFK LGPLSYQNMA FITAKDKYKL YPVRIPRLDT SKEFSAYVSG LFEIYRDLGD
DRVFNVPTIG VVNSNFAKEH NATVNLAMEA ILNELEVFIG RVKDQDGRVN RFYELEESLT
VLNCLRTMYF ILDGQDVEEN RSEFIESLLN WINRSDGEPD EEYIEQVFSV KDSTAGKKVF
ETQYFWKLLN QLVLRGLLSQ AIGCIERSDL LPYLSDTCAV SFDAVSDSIE LLKQYPKDSS

STFREWKNLV LKLSQAFGSS ATDISGELRD YIEDFLLVIG GNQRKILQYS RTWYESFCGF
LLYYIPSLEL SAEYLQMSLE ANVVDITNDW EQPCVDIISG KIHSILPVME SLDSCTAAFT
AMICEAKGLI ENIFEGEKNS DDYSNEDNEM LEDLFSYRNG MASYMLNSFA FELCSLGDKE
LWPVAIGLIA LSATGTRSAK KMVIAELLPH YPFVTNDDIE WMLSICVEWR LPEIAKEIYT
TLGNQMLSAH NIIESIANFS RAGKYELVKS YSWLLFEASC MEGQKLDDPV LNAIVSKNSP
AEDDVIIPQD ILDCVVTNSM RQTLAPYAVL SQFYELRDRE DWGQALRLLL LLIEFPYLPK
HYLVLLVAKF LYPIFLLDDK KLMDEDSVAT VIEVIETKWD DADEKSSNLY ETIIEADKSL
PSSMATLLKN LRKKLNFKLC QAFM

**Caldesmon**
MDDFERRREL RRQKREEMRL EAERLSYQRN DDDEEEAARE RRRRARQERL RQKEEGDVSG
EVTEKSEVNA QNSVAEEETK RSTDDEAALL ERLARREERR QKRLQEALER QKEFDPTITD
GSLSVPSRRE VNNVEENEIT GKEEKVETRQ GRCEIEETET VTKSYQRNNW RQDGEEEGKK
EEKDSEEEKP KEVPTEENQV DVAVEKSTDK EEVVETKTLA VNAENDTNAM LEGEQSITDA
ADKEKEEAEK EREKLEAEEK ERLKAEEEKK AAEEKQKAEE EKKAAEERER AKAEEEKRAA
EERERAKAEE ERKAAEERER AKAEEERKAA EERAKAEEER KAAEERAKAE EERKAAEERA
KAEKERKAAE ERERAKAEEE KRAAEEKARL EAEKLKEKKK MEEKKAQEEK AQANLLRKQE
EDKEAKVEAK KESLPEKLQP TSKKDQVKDN KDKEKAPKEE MKSVWDRKRG VPEQKAQNGE
RELTTPKLKS TENAFGRSNL KGAANAEAGS EKLKEKQQEA AVELDELKKR REERRKILEE
EEQKKKQEEA ERKIREEEEK KRMKEEIERR RAEAAEKRQK VPEDGVSEEK KPFKCFSPKG
SSLKIEERAE FLNKSAQKSG MKPAHTTAVV SKIDSRLEQY TSAVVGNKAA KPAKPAASDL
PVPAEGVRNI KSMWEKGNVF SSPGGTGTPN KETAGLKVGV SSRINEWLTK TPEGNKSPAP
KPSDLRPGDV SGKRNLWEKQ SVEKPAASSS KVTATGKKSE TNGLRQFEKE P

**Nup1pΔN**
IQESFVPNSE RSGTPTLKKN IEPKKDKESI VLPTVGFDFI KDNETPSKKT SPKATSSAGA
VFKSSVEMGK TDKSTKTAEA PTLSFNFSQK ANKTKAVDNT VPSTTLFNFG GKSDTVTSAS
QPFKFGKTSE KSENHTESDA PPKSTAPIFS FGKQEENGDE GDDENEPKRK RRLPVSEDTN
TKPLFDFGKT GDQKETKKGE SEKDASGKPS FVFGASDKQA EGTPLFTFGK KADVTSNIDS
SAQFTFGKAA TAKETHTKPS ETPATIVKKP TFTFGQSTSE NKISEGSAKP TFSFSKSEEE
RKSSPISNEA AKPSFSFPGK PVDVQAPTDD KTLKPTFSFT EPAQKDSSVV SEPKKPSFTF
ASSKTSQPKP LFSFGKSDAA KEPPGSNTSF SFTKPPANET DKRPTPPSFT FGGSTTNNTT
TTSTKPSFSF GAPESMKSTA STAAANTEKL SNGFSFTKFN HNKEKSNSPT SFFDGSASST
PIPVLGKPTD ATGNTTSKSA FSFGTANTNG TNASANSTSF SFNAPATGNG TTTTSNTSGT
NIAGTFNVGK PDQSIASGNT NGAGSAFGFS SSGTAATGAA SNQSSFNFGN NGAGGLNPFT
SATSSTNANA GLFNKPPSTN AQNVNVPSAF NFTGNNSTPG GGSVFNMNGN TNANTVFAGS
NNQPHQSQTP SFNTNSSFTP STVPNINFSG LNGGITNTAT NALRPSDIFG ANAASGSNSN
VTNPSSIFGG AGGVPTTSFG QPQSAPNQMG MGTNNGMSMG GGVMANRKIA RMRHSKR

**Fesselin**
MIQSAAPSIP RVEVILDCSD REKEAPKSLA ERGCVDSQVE GGQSEAPPSL PSFAISSEGT
EQGEDNQHSE KDHRPLKHRA RHARLRRSES LSEKQVKEAK SKCKSIALLL TAAPNPNSKG
VLMFKKRRQR ARKYTLVSYG TGELERDEDE GEEGEVEEGD KENTFEVSLL ATSESEIDED
FFSDIDNDKK IVTFDWDSGL LEVEKKTKSG DEMQTLPETT GKGALMFARR RQRMDQITAE
QEEMKARTAH AEEQREVTVS ENFQKVSSSA YQTKEEEMLR QQPCISKSYA DVSQNDGKIV
QQNGFGVAPD TSLSFQSSEA QKAASLNRTA KPFPFGVQNR AAAPFSPTRN VTSPLSDLPA
PPPYCSISPP PEALYRPLSA PAASKAAPIL WSHTEPTERI ASRDERIAVP AKRTGILQEA
KRRSTSKPMF SFKEAPKVSP NPALLSLVHN AEGKKGSGAG FESGPEEDYL SLGAEACNFM
QSQASKQKAP PPTAPKPSLK VSPAAGTPVS PVWSPAVASN KAPSFPAPAS PQAAYPAPLK
SPQYPHSPSA NPPNTLNLSG PFKGPQATLA SPNHPAKTPT TPSAGETKPF EMPPEMRGKG
AQLFARRHSR MEKYVVDSET VQANMARASS PTPSLPASWK YSSNVRAPPP VAYNPIHSPS
YPPAATKPFP KSTAATKNTK RKPKKGLNAL DIMKHQPYQL DASLFTFQPP SNKESLGIKQ
IPKLPTSKQA TSLRLPGSAS PTNVRASSVY SVPAYSSQPS FQSNASTPVN ESYTPTGYSA
FSKPESTTSS LFTAPRPKFS AKKAGVIAQE RSSGRSLSLP GKPSFISRAT SPTSPLIFQP
APDYFSKPDT AADKPGKRLT PWEAAAKSPL GLVDEAFRPQ NMQESIAANV VSAAHRKTLP
EPPDEWKQKV SYEPPGPSAS LALLGGKQPG VTSARKSSLS VSNATTQAGS QQQYAYCSQR
SQTDPDIMSM DSRSDYGLST ADSNYNPQPR GWRRPT
